# Multiplex genome editing of human pluripotent stem cells using Cpf1

**DOI:** 10.1101/2022.04.13.488123

**Authors:** Haiting Ma, Rudolf Jaenisch

**Affiliations:** Whitehead Institute for Biomedical Research, Cambridge, MA 02142, USA; Department of Biology, Massachusetts Institute of Technology, Cambridge, MA 02142

## Abstract

Targeted genome editing of human pluripotent stem cells (hPSC) is critical for basic and translational research, and can be achieved with site-specific endonucleases. Class 2 clustered regularly interspaced short palindromic repeats (CRISPR) Cpf1 represents a class of RNA programmable DNA endonucleases with features that could be complementary to the widely used Class 1 CRISPR-Cas9 system. Here we engineered a single vector system to deliver both Cpf1 and guide RNA, and achieved efficient knockin in transformed human cells. We then adopted the system to function in hPSCs by increasing expression levels of Cpf1, and showed that Cpf1 efficiently mediate homology directed repair in multiple loci (including knockin of a 6 kb cassette at the *INS*) in multiple hPSC lines, resulting in clones with transgenic cassettes integrated only at the targeted loci in the genome. Furthermore, with delivery of Cpf1 and a single U6 promoter-driven guide RNA array composed of an *AAVS1*-targeting guide and a *MAFB-*targeting guide, we demonstrated efficient multiplex genome editing in hPSCs at the *AAVS1* (knockin) and *MAFB* (knockout) loci. Targeted hPSCs expressed pluripotency markers, and could be directed to differentiate to pancreatic β-like cells, neural progenitor cells, and neurons *in vitro*. Generated *INS* reporter cells differentiated into β-like cells that express tdTomato and luciferase specifically in insulin-expressing cells, allowing for *in vivo* tracking of human β-like cells in humanized mice. By targeted screening of potential off-target sequences with most homology to guide RNA targeted sequences, we could not detect off-target mutations for all guide RNAs at the condition of efficient genome editing with Cpf1 in hPSCs. This work provides a system complementary to Cas9 for potentially precise genome editing in hPSCs.

## INTRODUCTION

With the development of human induced pluripotent stem cell (iPSC) technology (Takahashi et al., 2007; Wernig et al., 2007) and successful differentiation of functional cell types, autologous cell replacement therapies emerge as a potential treatments for multiple currently incurable diseases (Hochedlinger and Jaenisch, 2015). Genetic modifications of human pluripotent stem cells (hPSC), either correcting existing genetic lesions or conferring novel functions such as overexpressing PDL1 for increased immune tolerance (Ma et al., 2020), constitutes an indispensable prerequisite for realizing the full potential of hPSC therapies (Hockemeyer and Jaenisch, 2016). Several technologies have been developed for efficient genome editing in hPSCs: engineered zinc finger nucleases (Hockemeyer et al., 2009), TALE nucleases (Hockemeyer et al., 2011), and CRISPR-Cas9 (Cong et al., 2013; Mali et al., 2013). Complementary utilization of these systems is a critical component for hPSC-based regenerative medicine (Cornu et al., 2017; Cox et al., 2015).

The Class 1 RNA programmable CRISPR-Cas9 system enables efficient genome editing that can be conveniently applied to diverse targets *in vitro* and *in vivo* in a variety of cell types in multiple species (Hsu et al., 2014). However, wild type SpCas9 can generate double strand DNA breaks in off-target sites that are partially complementary to guide RNA. The off-target effects of SpCas9 can be reduced by modifying the basic amino acid residues of the DNA binding domain of SpCas9, but the modified SpCas9 could also show reduced activity compared to wild type SpCas9 (Slaymaker et al., 2016). Furthermore, to achieve effective multiplex editing at the single cells levels with DNA vector, each guide RNA requires its own promoter, thereby complicating multiplex applications (Sakuma et al., 2014). Moreover, the obligate 3’ nGG protospacer adjacent motif (PAM) sequences could limit its application in AT-rich targets.

CRISPR-Cpf1 family endonucleases were identified and shown to confer efficient genome editing in transformed mammalian cell lines (Zetsche et al., 2015), and in mouse embryos (Hur et al., 2016; Kim et al., 2016b). Cpf1 requires the 5’ T-rich PAM motif (Zetsche et al., 2015), therefore could be used for T-rich targets. In contrast to the blunt end double stranded DNA break products by Cas9, Cpf1 generates 5-nucleotid staggered cuts distal to the PAM sequences, and Cpf1 was proposed to improve homology directed repair (Zetsche et al., 2015). Additionally, *Francisella novicida* Cpf1 (FnCpf1) was shown to have RNase activity to process its cognate crRNA (Fonfara et al., 2016). Similar, *Acidaminococcus* Cpf1 (AsCpf1), a Cpf1 ortholog showing robust gene editing in mammalian cells (Zetsche et al., 2015), also harbors RNase activity sufficient for the maturation of its crRNA in cells, thereby making it possible to deliver multiple guide RNAs from a pre-crRNA array driven by a single U6 promoter (Zetsche et al., 2017). Furthermore, Cpf1 was shown to result in less off-target effects in genome editing experiments with transformed human cell lines (Kim et al., 2016a; Kleinstiver et al., 2016). Taken together, these unique features of Cpf1 could make it a system that complements the currently available gene-editing tools.

In this study, we developed a single vector system for the expression of AsCpf1 and its crRNA. Transfection of the Cpf1 vectors with its target crRNA resulted in robust mutations in targeted loci in HEK293T cells. Together with an *AAVS1* targeting Cpf1 construct, introduction of a donor construct resulted in knockin of *tdTomato* expression cassette at the *AAVS1* locus. Compared to SpCas9, we observed less efficient mutation formation and less efficient homology directed repair with Cpf1, suggesting mutagenesis efficiency is a major factor determining the efficiency of homology directed repair. We then utilized the system in hPSCs. In contrast to HEK293T cells, we could only detect mutation formation with AsCpf1 driven by the CAGGS promoter, not by the weaker CMV promoter that functioned in HEK293T cells. Consistently, with high expression levels of AsCpf1, we achieved efficient knockin of *tdTomato* expression cassette at the *AAVS1* locus in multiple hPSCs lines including iPS cells. Furthermore, by introducing a pre-crRNA array composed of *AAVS1* targeting guide, and *MAFB* targeting guide, we observed 100% editing of *MAFB* locus in *AAVS1* targeted clones, providing proof-of-principle example for genome editing in multiple loci in hPSCs at the single cell level. Finally, we generated insulin expression reporter clones through the knockin of a 6 kb cassette expressing both the *luciferase* and *tdTomato* at the *INS* locus in hPSCs, at an efficiency higher compared to TALE nuclease mediated knockin at the same locus, demonstrating the utility of Cpf1-mediated gene targeting in hPSCs.

## RESULTS

### A single plasmid system to deliver both AsCpf1 and guide RNA

To improve the system for genome editing including potential use for homology directed repair, we developed a bicistronic plasmid expressing AsCpf1 with an N-terminal SV40 nuclear localization signal (NLS) and a C-terminal nucleoplasmin NLS driven by the CMV promoter, and expressing AsCpf1 crRNA driven by human U6 promoter (**Figure 1A**). Guide RNA for target genes can be cloned with a pair of annealed oligonucleotides ligated to BbsI digested backbone vector (**Supplemental Figure 1, Figure 1**). Furthermore, to facilitate detection and fluorescence activated cell sorting (FACS) of transfected cells, we included a GFP coding cassette separated from the AsCpf1 open reading frame with a T2A sequence (**Figure 1A**). To test the single plasmid system, we designed an *AAVS1* target that lies close to a Cas9 site (Mali et al., 2013) that was cloned to a similar vector pX458 (Ran et al., 2013) that expressed SpCas9 separated from a GFP expressing cassette (**Figure 1A, Figure 1B** top panel). Compared to control sample (**Figure 1C**, CTL), we observed GFP signal in HEK293T cells transfected with plasmid backbone (**Figure 1C**, Cpf1-2A-GFP, no guide), and with the *AAVS1* guide (**Figure 1C** Cpf1-2A-GFP, *AAVS1* sg), similar to cells transfected with Cas9 expressing plasmid (**Figure 1C**, Cas9-2A-GFP no guide, Cas9-2A-GFP *AAVS1* sg). We extracted genomic DNA and performed T7EI assay, and observed cleavage of target regions into fragments with expected sizes from HEK293T cells transfected with the plasmid with the *AAVS1* guide RNA, whereas vector control did not show detectable editing of target sequences (**Figure 1D**). Moreover, expression of D908A mutant AsCpf1 that lacks nucleus activity (Yamano et al., 2016) and the same *AAVS1* guide RNA did not yield detectable mutation formation at the *AAVS1* locus (**Supplemental Figure 2**).

**Figure 1.**
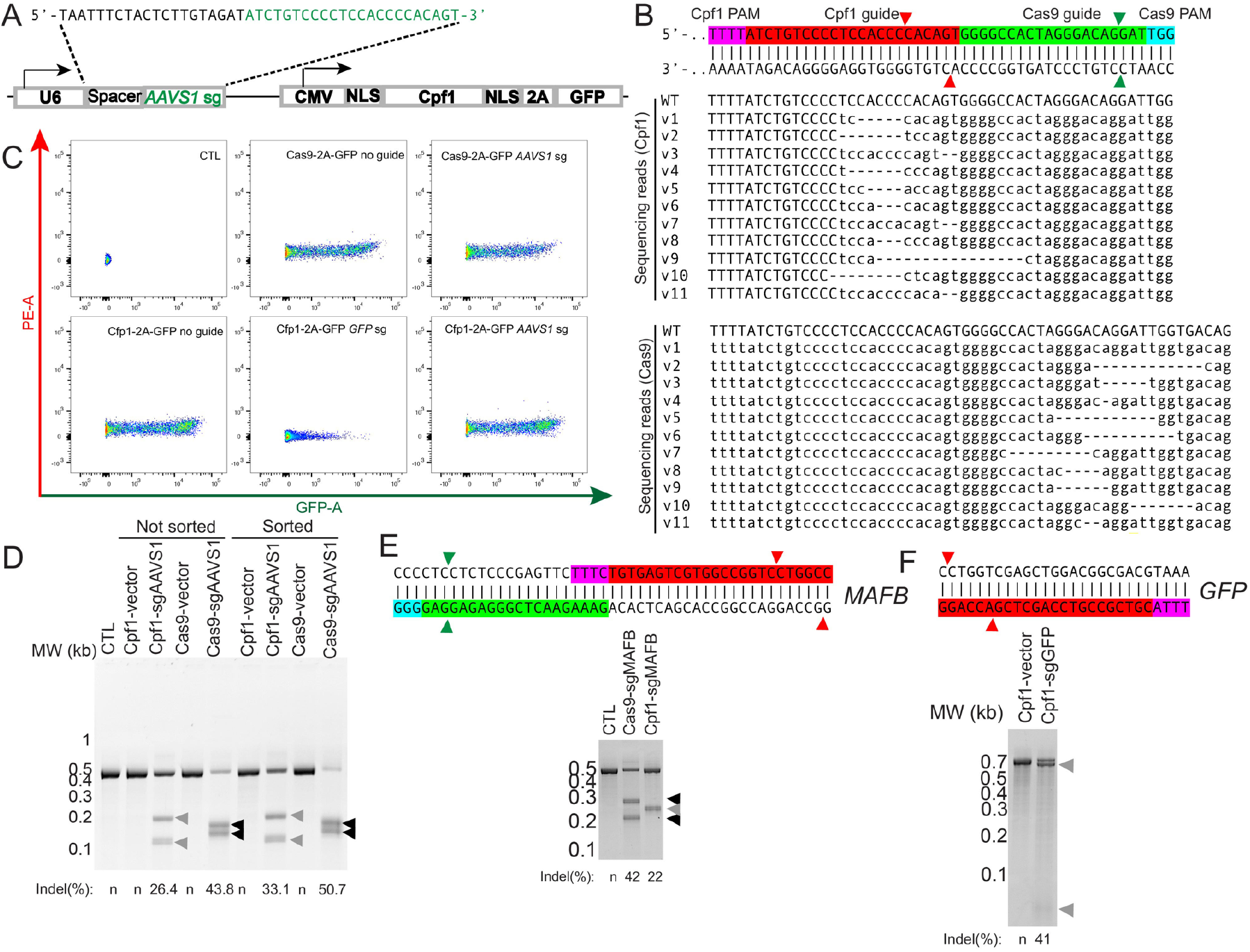
A single vector system for genome editing with AsCpf1. (A) Design of a bicistronic vector delivering both guide RNA (*AAVS1* sg RNA shown as nucleotides in green) and AsCpf1 followed by 2A-GFP. (B) Top panel: diagram showing the design of AsCpf1 *AAVS1* guide (in red) and SpCas9 *AAVS1* guide (in green). PAM sequences for AsCpf1 and SpCas9 are highlighted in magenta and cyan, respectively. AsCpf1 mediated double strand breaks are marked by red arrowheads, and SpCas9 mediated double strand breaks are marked by green arrowheads. Middle panel: sequencing results confirmed mutations in *AAVS1* locus caused by expression of Cpf1 together with its *AAVS1* guide. Bottom panel: sequencing results confirmed mutations upon expression of SpCas9 together with its *AAVS1* guide. (C)Representative FACS scatter plots of control HEK293T cells, SpCas9-2A-GFP expressing, SpCas9-2A-GFP *AAVS1* guide (green regions in top panel of E) expressing, AsCpf1-2A-GFP expressing, AsCpf1-2A-GFP *GFP* guide expressing, AsCpf1-2A-GFP *AAVS1* guide (red regions in top panel of (B) expressing HEK293T cells. (D) T7EI assay of mutations in the *AAVS1* locus from unsorted and sorted samples. Estimated on-target efficiencies are listed below the gel. (E) Top panel: a diagram showing the design of AsCpf1 *MAFB* guide (in red) and SpCas9 *MAFB* guide (in green). PAM sequences for AsCpf1 and SpCas9 are highlighted in magenta and cyan, respectively. AsCpf1 mediated double strand breaks are marked by red arrowheads, and SpCas9 mediated double strand breaks are marked by green arrowheads. Bottom panel: T7EI assay of mutations in the *MAFB* locus from sorted samples. Estimated on-target efficiencies are listed below the gel. (F) Top panel: a diagram showing the design of AsCpf1 *GFP* guide (in red). The PAM sequence for AsCpf1 is highlighted in magenta. AsCpf1 mediated double strand breaks are marked by red arrowheads. Bottom panel: T7EI assay of mutation in *GFP* gene with vector control, and *GFP* guide RNA expressing cells. Grey arrows denote expected T7EI digestion products. Estimated on-target efficiencies are listed below the gel.

To compare the on-target editing efficiency of AsCpf1 and SpCas9, we extracted genomic DNA from HEK293T cells transfected with vectors without guide RNAs, or plasmids with the corresponding *AAVS1* guide RNAs, performed T7EI assay, and observed cleavage of target regions into fragments with expected sizes (**Figure 1D**), with SpCas9 more effective than AsCpf1 (43.8% vs 26.4%). To compare results of cell populations that all expressed SpCas9 or AsCpf1, we sorted GFP positive cells with FACS, and performed T7EI analysis. As expected, sorted cells showed higher percentage of cleaved products compared to unsorted population (**Figure 1D**), and in the GFP positive populations, SpCas9 expressing cells showed higher cleavage efficiency (50.7%) compared to AsCpf1 (33.1%), demonstrating that Cas9 has higher on-targeting efficiency compared to AsCpf1 for these two guide RNAs.

To determine the sequences of AsCpf1 mediated genome editing products, we used next generation sequencing on the PCR amplicons from HEK293T cells transfected with *AAVS1* targeting AsCpf1 or SpCas9 plasmids, and identified multiple alleles in the *AAVS1* locus in HEK293T cells transfected with AsCpf1-*AAVS1* plasmid and SpCas9-*AAVS1* plasmid (**Figure 1B**, middle and bottom panel), with edited regions aligned to the expected double stranded DNA breaks generated by AsCpf1 (**Figure 1B**, red arrow heads in top panel), and SpCas9 (**Figure 1B**, green arrow heads in the top panel). In contrast, HEK293T cells transfected with no guide vector controls did not yield significant amounts of reads with mutations.

To test if this bicistronic AsCpf1 vector could generate mutations in other loci, we tested the editing efficacy of AsCpf1 and SpCas9 in *MAFB* (**Figure 1E**, top panel), and found similar results that SpCas9 is more efficient than AsCpf1 (42% vs 22%) (**Figure 1F**, bottom panel). To test if the bicistronic AsCpf1 system is effective for episomal targets, we generated a plasmid with a *GFP* guide Cpf1-2A-GFP GFP sg (**Figure 1F**). Compared to vector control, the expression level of GFP was lower in cells transfected with Cpf1-2A-GFP GFP sg (**Figure 1C** Cpf1-2A-GFP GFP sg). T7EI assay confirmed formation of mutations in the GFP gene upon Cpf1-2A-GFP GFP sg transfection (**Figure 1F**). Taken together, these data demonstrate that the single vector system allows AsCpf1-mediate mutation formation in mammalian cells upon cloning of guide RNAs.

### Homology directed repair by AsCpf1-mediated gene editing in transformed human cells

In contrast to Cas9, Cpf1 enzymes generate sticky ends that have been hypothesized to stimulate homology directed repair (Zetsche et al., 2015). We tested this by targeting the safe harbor *AAVS1* locus to generate site specific knockin. We modified an AAVS1-CAGGS-tdTomato gene trap donor vector (Addgene 159275) such that the PAM motifs for SpCas9 and AsCpf1 were both disrupted in the donor vector. Therefore, the same targeting construct was used as donor template for SpCas9 or AsCpf1 mediated homology directed repair mediated knockin (**Figure 2A**). We tested gene targeting in HEK293T cells facilitated by puromycin selection (**Figure 2A**, bottom panel), and obtained clones with ubiquitous expression of tdTomato with both SpCas9-*AAVS1*-sg and AsCpf1-*AAVS1*-sg co-transfected with the AAVS1-CAGGS-tdTomato donor vector (**Figure 2B**). To test correct knockin at the *AAVS1* locus rather than random integration, we designed a PCR based assay with primers specifically in the targeting vector (primer 489), and a primer outside of the homologous arm (primer 495) (**Figure 2A**). DNA samples from the cells with correct knockin gave rise to a PCR product of about 1.5 kb (**Figure 2C**, left panel). Sequencing of the PCR products confirmed the correct junction sequences from the endogenous gene that is not included in homologous arms and sequence in the homologous arm (**Figure 2C**, right panel). In contrast, control samples did not yield correct PCR products (**Figure 2C**). Quantification of the ratio of HEK293T cell clones with correct knockin based on the PCR assay to the total number of randomly picked clones showed that SpCas9 had slightly higher knockin efficiency compared to AsCpf1 mediated knockin, although the difference is not statistically significant (**Figure 2D**, P>0.05, non-paired t-test). These results demonstrate that AsCpf1 could be used to mediate homology directly repair in transformed human cells.

**Figure 2.**
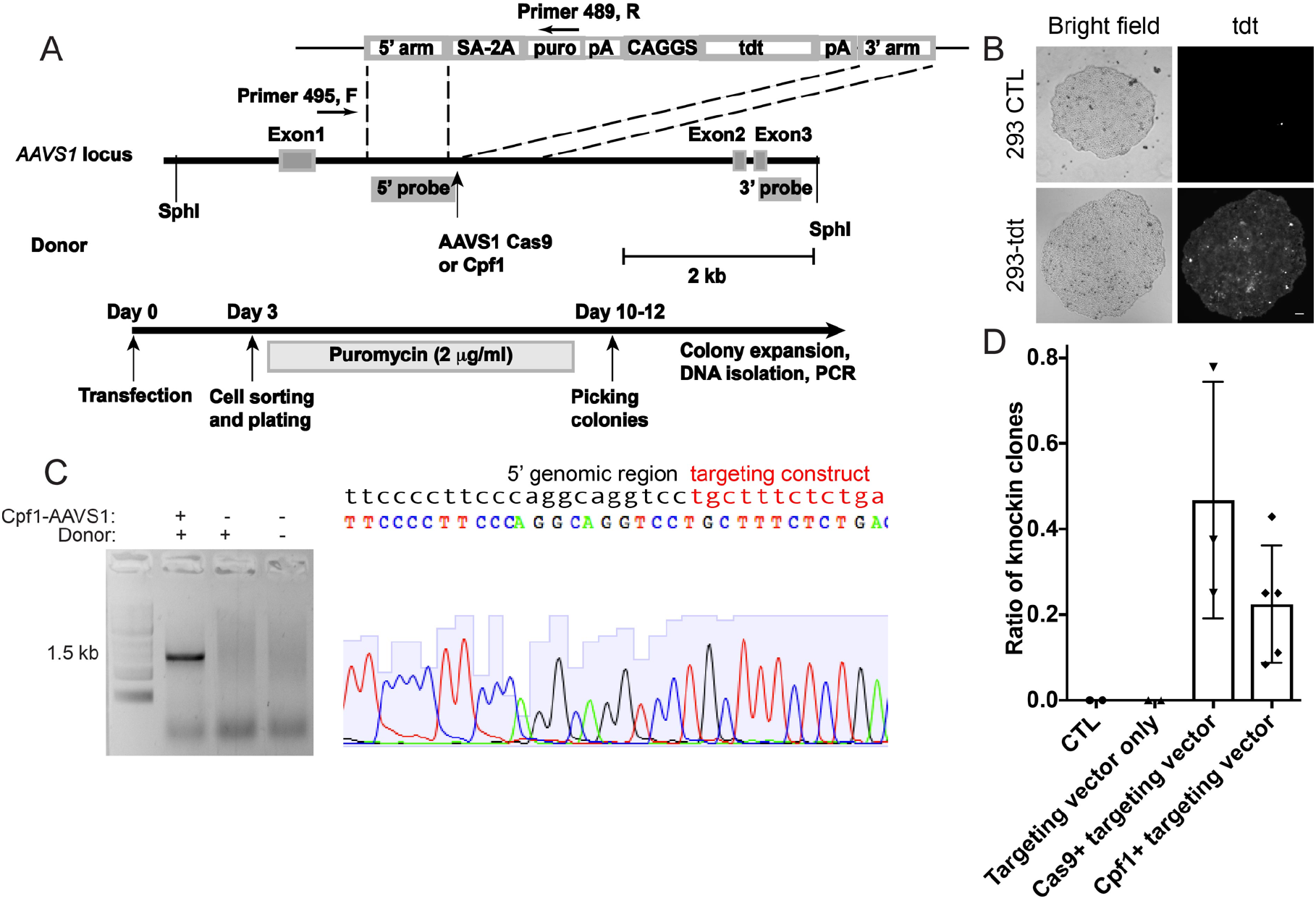
AsCpf1-mediated knockin of a tdTomato expressing cassette to the *AAVS1*locus in HEK293T cells. (A) A schematic illustrating targeting strategy with SpCas9 or AsCpf1 (top panel), and a timeline to generate HEK293T cells with CAGGS-tdTomato knockin at the *AAVS1* locus (bottom panel). (B) Expression of tdTomato in AsCpf1 targeted HEK293T cells (bottom panel), but not in the control cells (top panel). (C) Left panel: representative genotyping PCR results with primer 495 and 489 in (A) to identify HEK293T cell clones with the CAGGS-tdTomato cassette integrated at the *AAVS1* locus. Right panel: a representative Sanger sequencing chromatogram by sequencing the junction region of the PCR amplicon from targeted cells. Sequences inside and outside of the homology arm are shown in red and black, respectively. (D) Knockin efficiencies (ratio of randomly picked clones that show positive genotyping PCR results as in C) mediated by SpCas9 or AsCpf1 are plotted.

### Cpf1-mediated genome editing in hPSCs with a CAGGS promoter driven AsCpf1

We then tested whether AsCpf1 could mediate homology directed repair in hPSCs by targeting hPSC cell line H1 (Thomson et al., 1998) with SpCas9 and AsCpf1 plasmids targeting *AAVS1* (**Figure 1B**) and the AAVS1-CAGGS-tdTomato donor template (**Figure 2A**). From SpCas9 targeted experiments, all of the randomly picked colonies showed correct targeting by PCR assay (**Supplemental Figure 3**) and can identify correct targeted clones by Southern blot. In contrast, none of the randomly picked colonies from AsCpf1 targeted cells showed correct PCR results or positive results with Southern blot (**Supplemental Figure 3**), indicating the colonies arose from integration of targeting vector into loci other than the *AAVS1* locus the genome.

We observed strong GFP expression signals in HEK293T cells transfected with the bicistronic plasmid with CMV driven AsCpf1-2A-GFP (**Figure 1C**). However, in hPSCs, the expression levels of GFP were very low with the CMV promoter (**Figure 3A**, right side, top panel). To improve the expression level of AsCpf1, we generated a construct similar to the design in **Figure 1A**, with the replacement of CMV promoter by CAGGS promoter (**Figure 3A**, left panel). Co-electroporation of this CAGGS-AsCpf1-2A-GFP plasmid resulted in improvement of expression of GFP (**Figure 3A**, right side, bottom panel), suggesting expression of AsCpf1 was also increased.

**Figure 3.**
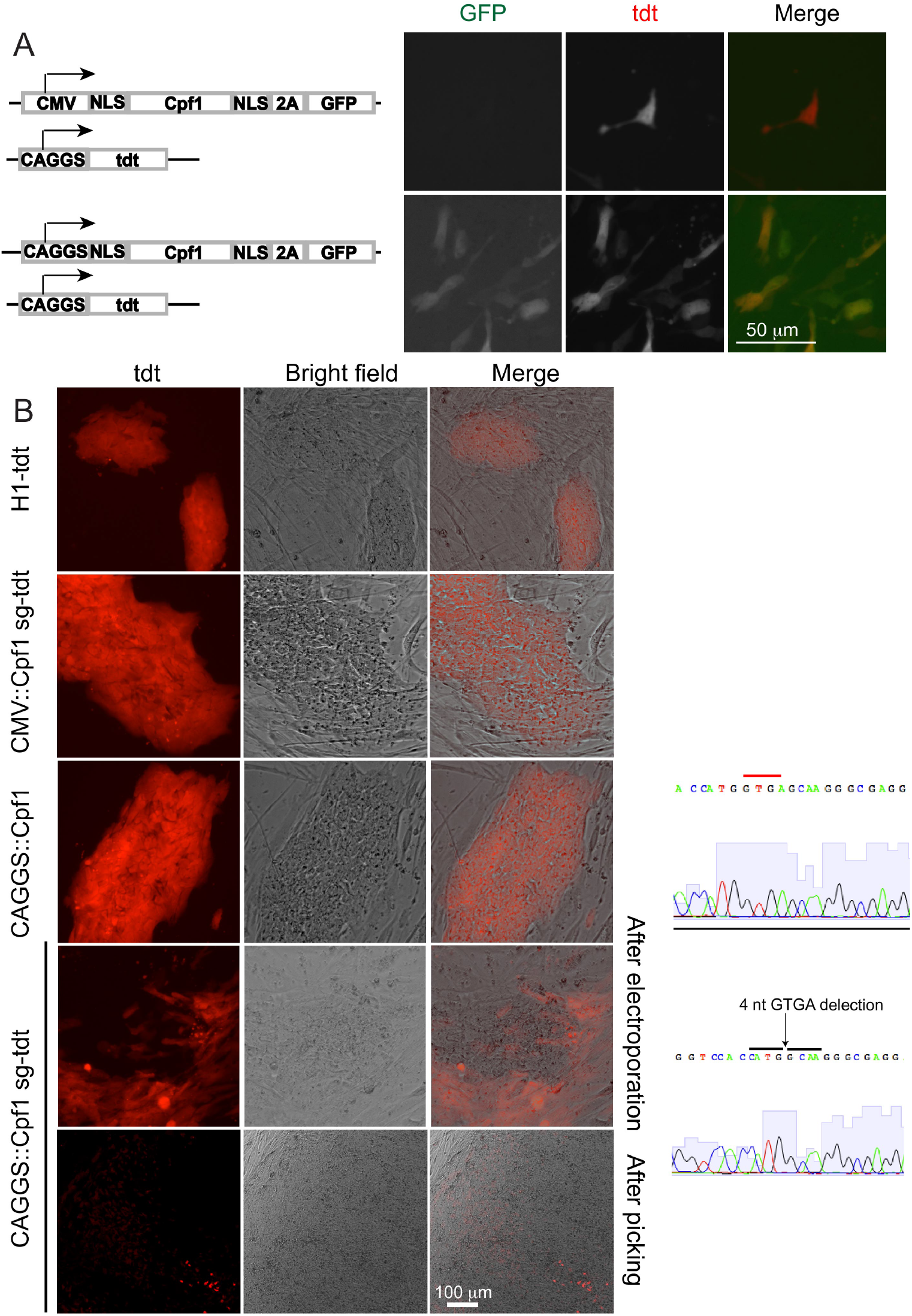
AsCpf1 mediated mutation formation in hPSCs. (A) Representative fluorescent micrographs of hPSCs electroporated with the corresponding constructs on the left. (B) Disruption of tdTomato expression in H1-tdT cells by CAGGS promoter driven AsCpf1 with a tdTomoto guide RNA, but not by the CMV promoter construct with the same guide RNA. A 4-nucleotide deletion was identified in cells that lost tdTomato expression, but not in control cells.

We tested the CAGGS-AsCpf1 construct by introducing a guide RNA targeting the tdTomato coding sequence (**Supplemental Figure 4A**, top panel), and transfected HEK293T cells that stably express tdTomato (HEK293-tdt) (**Figure 2B**). As expected, we observed expression of GFP, and increased tdTomato-low and negative cell proportion in the GFP positive population compared to control cells (**Supplemental Figure 4B**). At 4 days post transfection, tdTomato low cells can be observed in the HEK293T-tdTomato cells, particularly in cells with high expression of GFP (**Supplemental Figure 4A**, bottom panel). The loss of expression of tdTomato became clearer after propagation of sorted cells with low tdTomato expression (**Supplemental Figure 4C**, left panel). To test if the loss of tdTomato expression in cells was caused by mutations of the tdTomato coding sequence, we amplified the potentially mutated regions in tdTomato coding sequence with PCR, and observed out-of-frame mutation, consistent with loss-of-function of the tdTomato gene by premature stop codon (**Supplemental Figure 4C**, right panel). In contrast, the mutation could not be observed in parental HEK293 tdTomato cells or the cells expressing CAGGS-AsCpf1-2A-GFP vector. These results indicate that the CAGGS-AsCpf1-2A-GFP vector system could generate mutations in HEK293T cells.

Next, we electroporated the CAGGS-AsCpf1-2A-GFP construct with tdTomato guides to H1 cells with heterozygous integration of CAGGS-tdTomato expression cassette at the *AAVS1* locus (Ma et al., 2018), and observed increased expression of GFP with FACS analysis. We plated cells sorted for high expression of GFP, and detected tdTomato-negative clones (**Figure 3B**). In contrast, experiments performed using CMV-AsCpf1-2A-GFP sg-tdt, or CAGGS-AsCpf1-2A-GFP control did not resulted in tdTomato-negative clones (**Figure 3B**). We analyzed the tdTomato coding sequencing from the tdTomato-negative clones, and observed disruption of the tdTomato gene (4-nucleotide deletion) at the expected region edited by AsCpf1 with the tdTomato guide RNA (**Figure 3B**). In contrast, sequencing results from H1-tdTomato cells electroporated with control constructs did not yield detectable mutation in the tdTomato coding sequence (**Figure 3B**). These results indicate that increasing expression levels of Cpf1 by CAGGS promoter could more efficiently generate mutations in hPSCs than the CMV promoter driven AsCpf1.

### AsCpf1 mediated knockin at the *AAVS1* locus of hPSCs

We next tested the possibility of Cpf1-mediate knockin in hPSCs by introducing both the *AAVS1*-targeting CAGGS-Cpf1 construct and the AAVS1-CAGGS-tdTomato donor vector (**Figure 2A**) in WIBR3 cells (Lengner et al., 2010) (**Figure 4A**). With low amounts of *AAVS1*-targeting CAGGS-Cpf1 plasmid, we could not detect pluripotent stem cell clones showing positive PCR results indicating correct knockin at the *AAVS1* locus (**Table 1**). As we increase the amounts of *AAVS1*-targeting CAGGS-Cpf1 construct to 100 μg, we observed clones with uniform expression of tdTomato (**Figure 4B**). We isolated DNA from tdTomato positive clones and tested with Southern blotting with the probes used previously (Ma et al., 2020). Although we observed ectopic copy of the integrated CAGGS-tdTomato cassette (**Figure 4C**, clone 6), and clones showing integrated sizes different from expected size (**Figure 4C**, clone 5, and clone 7), 4 of 8 randomly picked clones were heterozygous with the CAGGS-tdTomato expression cassette integrated only at the *AAVS1* locus based on Southern blot analysis (**Figure 4C, Table 1**). Consistently, PCR tests and sequencing PCR amplicons showed correct junction sequence. To test if the targeted clones expressed pluripotency markers, we performed immunofluorescence, and detected expression of OCT4, NANOG, TRA-1-60, and SSEA4 similar to untargeted parental WIBR3 cells (**Figure 4D**). Additionally, the targeted cells generated teratoma composed of three germs layers upon transplantation in immunocompromised mice (**Figure 4I**, top panel), suggesting proper knockin in human pluripotent stem cells by Cpf1 did not interfere with pluripotency of targeted cells. We also sequenced the non-targeted *AAVS1* allele, and found 2 of 4 clones showed intact sequence for the non-targeted *AAVS1* allele, and the other two showed mutation in the non-targeted *AAVS1* allele. Moreover, we could generate CAGGS-tdTomato knockin at the *AAVS1* locus in iPS cells from a Niemann-Pick disease type C (NPC) patient (Maetzel et al., 2014) (**Table 1, Supplemental Figure 5**). These data demonstrate Cpf1 driven by CAGGS promoter could mediate knockin at the *AAVS1* locus in hPSCs with about 50% targeting efficiency (**Table 1**).

**Table 1.**
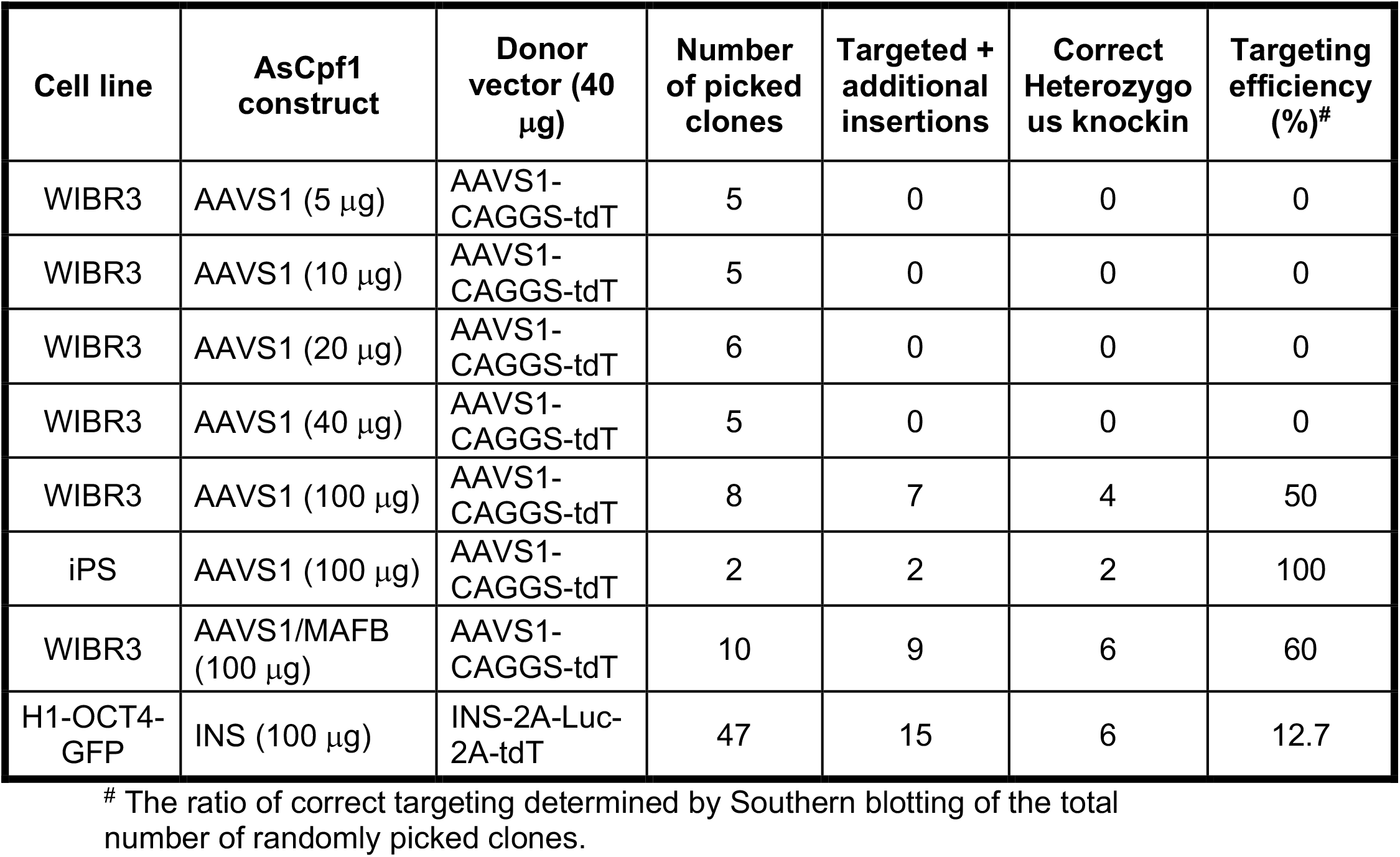
Summary of targeting efficiency in hPSCs.

**Figure 4.**
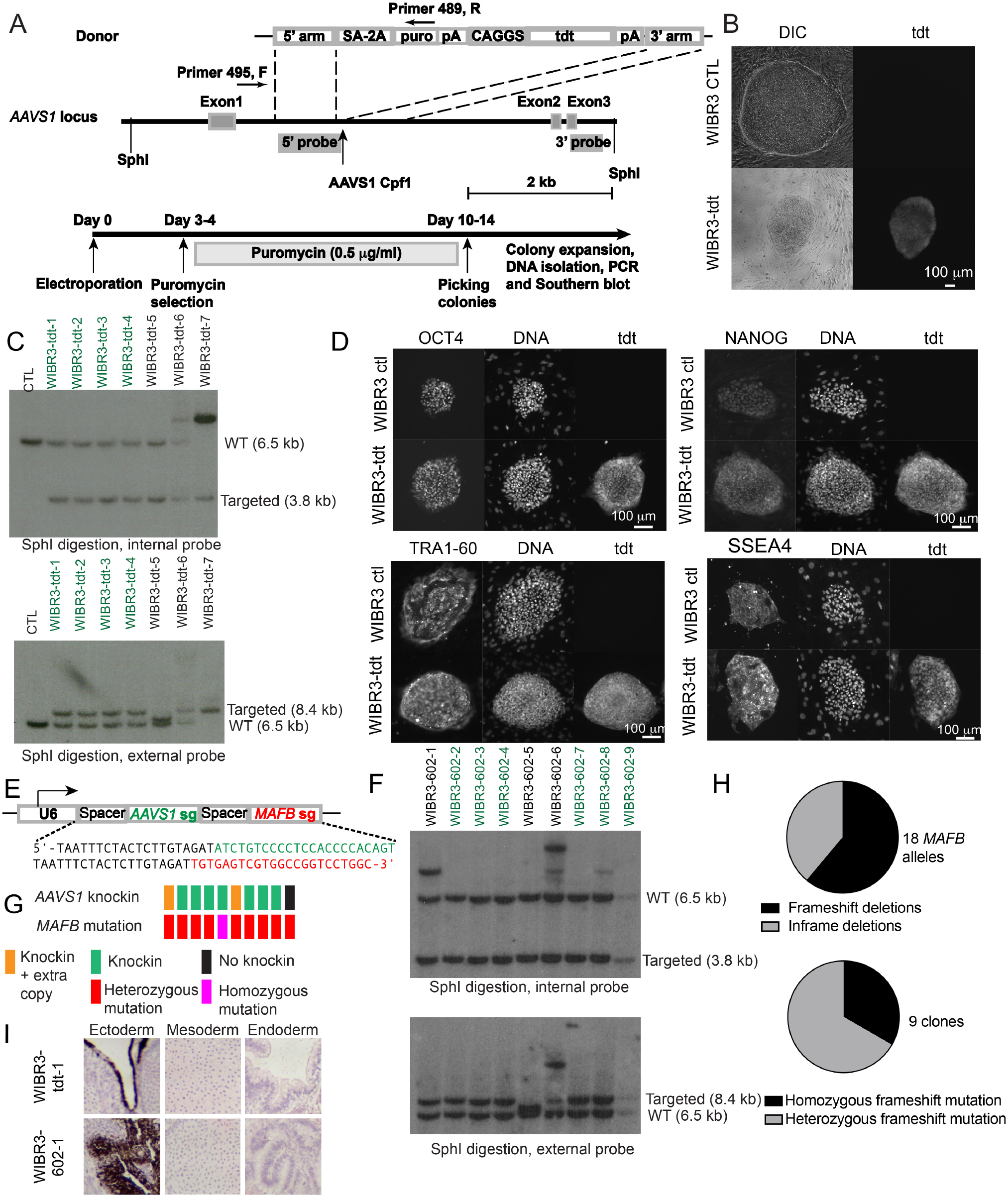
Multiplex genome editing in hPSCs with AsCpf1. (A) A schematics illustrating strategy for knockin at the *AAVS1* locus in hPSCs with Cpf1 (top panel), and timeline for the experiments (bottom panel). (B) Expression of tdTomato in a representative knockin clone but not in control. (C) Verification of correct knockin with Southern blot with internal probe (top panel) and external probe (bottom panel). Clones showing correct knockin are labeled in green. (D) Expression of pluripotency markers OCT4, NANOG, TRA1-60, and SSEA4 in targeted clones. (E) The design of an *AAVS1* and *MAFB* targeting spacer guide RNA array. (F) Southern blotting analyses of knocking of the *AAVS1* locus with the guide RNA array shown in (E). (G) A diagram showing the distribution of knockin and *MAFB* mutations in isolated hPSC clones. Each vertical bar pair represents an isolated hPSC clone. (H) Distribution of different mutation alleles in the *MAFB* gene upon AsCpf1 mediated editing with the guide array in (E). (I) H&E staining of teratoma tissues formed upon injection of WIBR3-tdTomato cells (top panels) or WIBR3-602-1 cells into immunocompromised mice.

### Simultaneous knockout of *MAFB* and knockin at *AAVS1* in hPSCs

Cpf1-mediated multiplex gene editing can be achieved with a single U6 promoter driven guide RNA array composed of multiple guide RNA targeting different loci (Zetsche et al., 2017). We tested multiplex editing by assembling a guide RNA array by combining the *AAVS1* guide and *MAFB* guide into the CAGGS-Cpf1-2A-GFP vector (construct 602) (**Figure 4E**). Upon transfection to HEK293T cells with CAGGS-Cpf1-2A-GFP-AAVS1-MAFB guide plasmid, we detected mutation formation at both the *AAVS1* and the *MAFB* loci (**Supplemental Figure 6**). We then introduced the CAGGS-Cpf1-2A-GFP-AAVS1-MAFB guide plasmid and the AAVS1-CAGGS-tdTomato donor vector to WIBR3 cells. After puromycin selection, in 10 randomly picked clones, we identified that 9 clones showed *AAVS1* knockin with PCR assay (**Supplemental Figure 7** top panel, **Table 1**) that showed expected junction sequence upon sequencing (**Supplemental Figure 7** bottom panel). Proper knockin at the *AAVS1* locus with the guide RNA array was confirmed by Southern blot analysis (**Figure 4F**). We also observed clones showing ectopic insertions (**Figure 4F**, clone 1 and clone 6), and abnormal band sizes (**Figure 4F**, clone 5). The targeting efficiency (60%) was similar to that with *AAVS1* guide only (**Table 1**). We also amplified the *MAFB* region containing the Cpf1 MAFB target (**Supplemental Figure 7** middle panel), and observed MAFB amplicons suggesting mutation formation in targeted clones but not in controls, indicating mutation mediated by the CAGGS-Cpf1-2A-GFP-AAVS1-MAFB expression. With sequencing of the MAFB amplicon, we found 10/10 clones showed mutations at the MAFB locus (**Figure 4G**), whereas parental control did not. Consistent with the high biallelic editing efficiency of the *MAFB* locus, we observed that 5 of 6 clones showing correctly targeting at the *AAVS1* locus also harbored mutations for the non-targeted *AAVS1* allele. We further analyzed *MAFB* mutation alleles of the 9 clones with *AAVS1* knockin. Most MAFB mutant alleles (11 out of 18) showed out-of-frame deletions (**Figure 4H**, top panel), resulting in 3 of 9 clones harbor out-of-frame deletions at both alleles (**Figure 4H**, bottom panel). These cells also form teratoma tissues in immunocompromised mice (**Figure 4I**, bottom panel), similar to targeted WIBR3-tdTomato cells, suggesting pluripotency is not affected with multiplex targeting. These results demonstrate robust multiplex genome editing in hPSCs with a Cpf1 guide RNA array driven by a single U6 promoter, and the possibility of generating hPSC loss-of-function alleles with simultaneous knockin at a different locus in a single experiment.

### Differentiation of AsCpf1-edited hPSCs to functional cell types

To evaluate whether AsCpf1-mediated genome editing could be used for modifying genes not expressed in hPSCs to generate reporter alleles, we used an *INS* gene targeting Cpf1 sgRNA (Ma et al., 2020) (**Figure 5A**). We used the puromycin resistant INS-2a-luciferase-2a-tdTomato-CAGGS-Puro donor construct (Ma et al., 2020) that, upon correct targeting, results in the expression of luciferase and tdTomato in insulin expressing β-like cells (**Figure 5B**). We then targeted H1 OCT4-GPF pluripotent stem cells (Zwaka and Thomson, 2003) for the generation of insulin luciferase-2A-tdt reporter (**Figure 5**). With PCR assay of 47 randomly selected puromycin resistant clones, 15 were positive. Southern blotting analysis showed 6 clones were correctly targeted with a heterozygous integration confirmed by Southern blot (**Figure 5C**). The targeted clones were tdTomato negative at the pluripotent stem cell state. We differentiated the targeted *INS* reporter hPSCs toward β-like cells with a published protocol (Ma et al., 2020), and identified detectable tdTomato signal at day 2 of stage 5 (**Figure 5D**, S5D2), and more tdTomato expressing cells were detected during later stages (**Figure 5D**). Immunofluorescence analyses showed that C-peptide expression cells also expressed tdTomato (**Figure 5E**). Quantitative RT-PCR analysis of cells sorted based on tdTomato expression levels showed high levels of *INS* and *luciferase* expression in tdTomato positive cells compared to tdTomato negative or lowly expressing cells (**Figure 5F**). This is consistent with specific detection of luciferase signal *in vitro* differentiated β-like cells but not in undifferentiated pluripotent stem cells (**Figure 5G**). Additionally, we transplanted the luciferase expressing β-like cells to the kidney capsule of immunocompromised mice, and could detect presence of *INS* expressing human β-like cells with *in vivo* imaging after surgical recovery (**Figure 5H**). Furthermore, human insulin was detected in blood circulation of mice showing luciferase signals, but not in control mice that did not show luciferase signal (**Figure 5H**). Additionally, we differentiated Cpf1 edited WIBR3-tdT cells to neural precursors as well as neurons, and obtained similar morphology and neuronal marker expression compared to control WIBR3 cells (**Supplemental Figure 8**). Taken together, these data suggest that Cpf1 could be used to target genes not expressed in hPSCs, and the targeted cells could be properly differentiated into functional cell types such as β-like cells and neurons.

**Figure 5.**
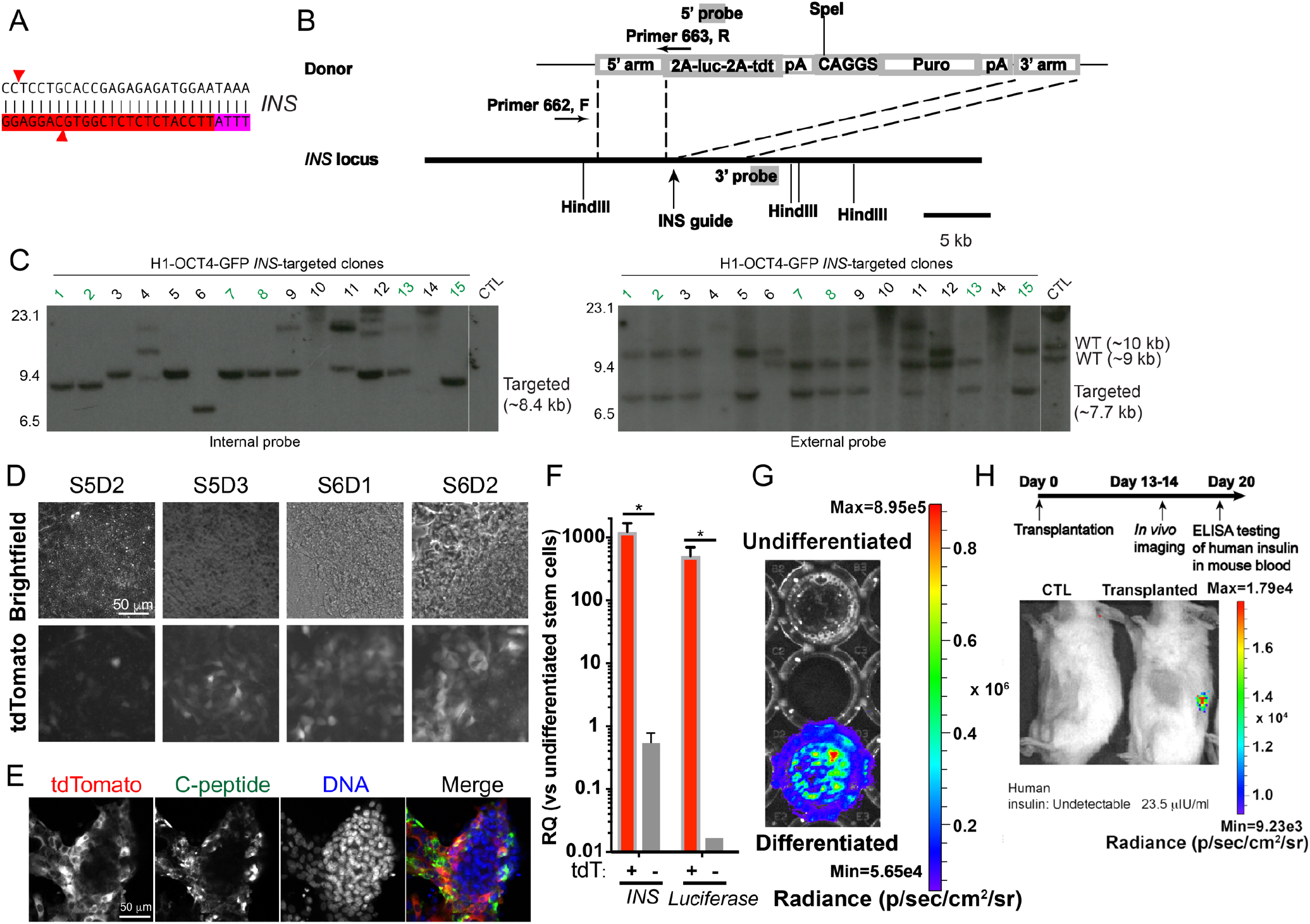
AsCpf1-mediated targeting in the *INS* locus in hPSCs enables monitoring of INS expression *in vitro* and *in vivo*. (A) The design of an *INS* targeting guide RNA. PAM sequence is highlighted in magenta, guide sequence is highlighted in red, and expected double stranded DNA breaks are marked by the red arrowheads. (B) The targeting strategy for generating *INS* luciferase and tdTomato reporter *INS-luciferase*-*tdT* allele by knocking 2A-luciferase-2A-tdt cassette to the 3’ end of the *INS* coding sequence. (C) Southern blots with internal probe (left panel) and external probe (right panel) to identify clones with a single copy of 2A-luciferase-2A-tdt cassette inserted into the *INS* locus. Correct clones were labeled in green. The double bands in the control samples for the external probe (right panels) correspond to two wild-type alleles of the H1-OCT4-GFP cells. (D) Emergence of tdTomato-expressing cells during the differentiation of the *INS-luciferase*-*tdT*reporter hPSCs toward pancreatic β-like cells during the differentiation. (E) Representative confocal micrographs of *INS-luciferase*-*tdT* hPSC differentiated β-like cells with anti-tdTomato staining (red) and anti-C-peptide staining (green). (F) Quantitative RT-PCR analysis of sorted tdTomato positive cells showed significantly higher expression of INS and luciferase compared to sorted tdTomato negative cells (* P<0.05, n=3 experiments). (G) β-like cells specific luciferase signal at stage 6 day 7. (H) Top panel: experimental design. Bottom panel: live mice luciferase imaging showed that β-like cells specific luciferase signals were detected in mice transplanted with β-like cells differentiated from *INS-luciferase*-*tdT* reporter hPSCs, but not in control, and luciferase signals are consistent with the presence of human insulin in mouse blood circulation.

### Evaluation of off-target effects in hPSC clones targeted with AsCpf1

Off-target effects can be generated by programmable nucleases through non-homologous end joining (NHEJ) mechanisms, and they constitute a major limitation for the application of these genome editing tools (Cornu et al., 2017; Cox et al., 2015). AsCpf1 was shown to have reduced off-target effects (Kim et al., 2016a; Kleinstiver et al., 2016) based on experiments with transformed human cell lines. We evaluated off-target effects in the hPSC clones with correct knockin by searching for sequences in the human genome that share most homology to the target sequences and are flanked by TTTn PAM motifs with Cas-OFFinder (Bae et al., 2014). For *AAVS1, MAFB*, and *INS* guide sequences used in this study, no sites with fewer than 4 mismatches were identified in the human genome. The number of sequences with 4 mismatched to *AAVS1, MAFB*, and *INS* guides are 5, 2, and 1, respectively (**Table 2**). We focused on the sequences with the least number of mismatches, and tested by PCR amplification of the potential off-target cutting site with genome DNA from PSC clones properly targeted by the corresponding guides, and sequencing PCR amplicons. No detectable off-target effects were identified in these sites with 4 mismatches for any of the Cpf1 guides used in generating knockin alleles in edited hPSC clones (**Table 2**). These results suggest that the system we developed for genome engineering Cpf1 could be used to generate targeted hPSC clones that are likely to have low levels of NHEJ mediated mutations at potential off-target sequences.

**Table 2.**
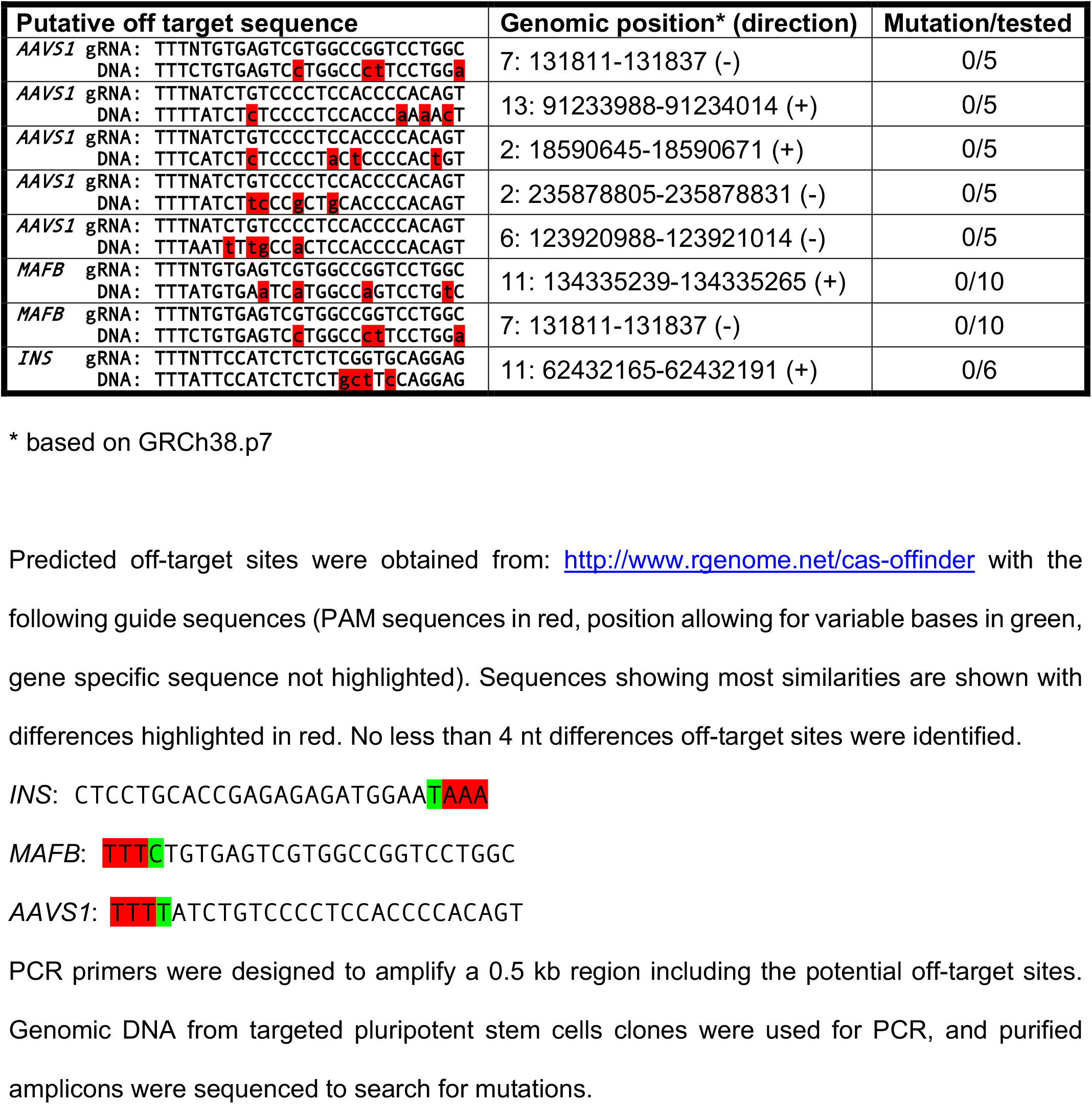
Experimental evaluation of off targeting effects of AsCpf1-mediated genome editing in hPSCs.

## DISCUSSION

In this study, we aimed to adopt the Cpf1 as a tool for genome editing, particularly precise genome editing via homology directed repair mediate locus specific knockin in hPSCs. Toward this goal, we developed a single vector system to deliver both AsCpf1 and guide sequences that could be cloned after the guide RNA scaffold. We achieved knockin at the *AAVS1* locus in transformed human cells. Through improving expression levels by using a CAGGS promoter and increasing the amount of Cpf1 plasmid electroporated to hPSCs, we were able to generate mutations in multiple genes in diverse hPSC cell lines including iPS cells. Furthermore, we could target mouse ES cells with Cpf1 (data not shown). Based on the capability of AsCpf1 to process its guide RNA through RNase activity, we were able to show multiplex genome editing (simultaneous targeting of *MAFB* (out-of-frame mutations) and *AAVS1* (knockin)) in hPSCs with a guide RNA array driven by a single U6 promoter, consistent with previous report in transformed human cells and in mouse *in vivo* (Zetsche et al., 2017). Human PSC edited with AsCpf1 could be differentiated to C-peptide positive β-like cells and neurons. Finally, we evaluated the potential off-target effects of Cpf1 in targeted hPSCs by analyzing loci that have 4-nucleotide difference (most similar to guide RNA sequences) from *AAVS1, MAFB*, and *INS* guide RNAs, and could not detect any alteration in all the clones we examined, indicating that the condition of gene generating with AsCpf1 could be used for precise genome editing in hPSCs.

With the single vector system that allows for sorting of cells expressing both AsCpf1 and guide RNA (**Figure 1**), we were able to compare on-targeting editing efficiency between SpCas9 in a similar system (Ran et al., 2013). In the loci examined, we observed more efficient on-targeting editing efficiency in SpCas9 compared to AsCpf1 in human cells (**Figure 1**), as well in mouse ES cells (data not shown). Consistently, we observed higher knockin efficiency mediated by SpCas9 compared to AsCpf1 (**Figure 2**). These results suggest that the on-target editing efficiency is a critical determinant for efficient homologous recombination. Furthermore, we did not observe a dramatic increment of homologous recombination efficiency ascribable to the 5’ overhang at the nascent DNA double stranded breaks generated by AsCpf1. This is also consistent with the experiments with zinc finger proteins and TALE proteins that are usually fused with FokI enzymes that generate 5’ overhang. Further studies are needed to investigate the role of DNA overhang in stimulating homology directed repair.

With AsCpf1-mediated targeting at the *AAVS1* locus, similar efficiencies (**Table 1**) were achieved compared with zinc finger nuclease mediated targeting (Hockemeyer et al., 2009). While in zinc finger nuclease targeted heterozygous *AAVS1* knockin clones, the untargeted *AAVS1* allele is generally wild-type (data not shown), we observed higher (50%-83%) editing efficiency in the non-targeted *AAVS1* allele in clones showing correct heterozygous knockin. In contrast, in the clones with heterozygous knockin at the *INS* locus with Cpf1 guide (**Figure 5A**), we did not observe alteration of sequences of the other *INS* allele, possibly because the on-target editing efficiency is lowers for the *INS* guide (**Figure 5A**) than the *AAVS1* guide (**Figure 1D**). With this *INS* Cpf1 guide, and a similar targeting construct with longer insertion cassette, we observed improved targeting efficiency (12.7%) at the *INS* locus compared to TALE nuclease-mediated knockin (3.1%) (Liu et al., 2014).

Cpf1 mediated gene disruption was achieved in mice by pronuclear injection of high concentration of mRNA and guide (Watkins-Chow et al., 2017). In this study, we also noted that expression of Cpf1 with a strong promoter (CAGGS), and high dose of Cpf1 (100 μg Cpf1 plasmid) were needed for efficient genome editing in hPSCs. Importantly, in clones correctely targeted with Cpf1, we did not detect off-target effects in loci with highest homologies to guide RNA sequences. These results are consistent with previous investigations on specificity of Cpf1 (Kim et al., 2016a; Kleinstiver et al., 2016). Based on the unique property of Cpf1 to perform multiplex genome editing with a guide RNA array driven by a single U6 promoter (Zetsche et al., 2017), we achieved high efficiency in multiple editing in hPSCs (**Figure 4G**). Taken together, this study demonstrates the AsCpf1-mediated genome editing system as a component of the genome editing tools for hPSCs.

## MATERIALS AND METHODS

### Cloning

PCR fragment with pY010 (Addgene #69982) as template and the following primers was cloned into pX458 (Addgene # 48138) at the AgeI FseI sites.

Forward:

GGACCGGTGGCCGCCACCATGGCCCCAAAGAAGAAGCGGAAGGTCGGTATCCACGGAGTCCCAGCAGCCACACAGTTCGAGGGCTTTACC

Reverse: TGCCTGGCCGGCCTTTTTCGT

Then the annealed product of

Forward: CACCGTAATTTCTACTCTTGTAGATGTCTTCGAGAAGACTTTTTTTT

Reverse: AAACAAAAAAAAGTCTTCTCGAAGACATCTACAAGAGTAGAAATTAC was cloned into the above plasmid at BbsI sites, giving rise to p-63.

Annealed products of gene specific oligo pairs (**Supplemental Table 1**) were cloned into the BbsI sites of p-63 to generate AsCpf1 with gene specific guide RNA.

p-69 was generated by cloning the PmlI and FseI fragment of a D908A (GAT → GcT) AsCpf1 to CAGGS:AsCpf1-2A-GFP (Addgene 159281).

### Transfection of HEK293T cells, FACS, and T7EI assay

HEK293T cells were cultured in DMEM high glucose medium (Gibco) supplemented with 10% FBS, 1X L-glutamine, 1X MEM-NEAA, and 1X penicillin/streptomycin (Life Technologies). Upon 60-80% confluence, cells cultured in 12-well plates (Corning) were transfected with 0.5 μg AsCpf1 or SpCas9 plasmid using Fugene 6 transfection reagent (Promega) according to manufacturer’s guideline. Three days posted transfection, cells were dissociated with trypsin, and GFP positive cells were sorted on a FACSAria (BD Biosciences), followed by DNA purification with DNeasy Blood & Tissue Kit (Qiagen).

PCR reactions were performed with 0.5 μg genomic DNA as template, and PCR with AmpliTaq Gold DNA Polymerase (Life technology) with the following PCR condition: 56**°**C, and 72**°**C extension for 50 seconds for 35 cycles, and extension at 72**°**C for 5 min in the end. PCR products were purified with PCR purification kit (Omega Biotech). A total of 200 ng PCR products were annealed in NEB buffer 2, and processed with 5 units of T7EI (NEB) for 1 hour at 37 **°**C before resolved on 10% TBE PAGE gels (BioRad) run at 100 volts for 2 hours. DNA fragments were observed with staining with ethidium bromide, and images were taken with an AlphaImager gel documentation system (Alpha Innotech). Mutation efficiencies were measured as previously reported (Ran et al., 2013).

### NGS sequencing of PCR products

PCR reactions with AmpliTaq Gold DNA Polymerase (Life technology) were set up with 0.5 μg genomic DNA and primer 483-484 (Supplemental Table 1) (anneal at 57**°**C, and amplified for 30 sec for 35 cycles). Amplicons were resolved on a 1.5% agarose gel followed by gel purification (Omega Biotech). Purified PCR products were dissolved in 35 microliter 10 mM Tris buffer (pH 8.0) at a concentration of 10-20 ng/μl. The ensuing NGS and data analysis was performed by the CCIB DNA Core Facility at Massachusetts General Hospital (Cambridge, MA). Briefly, partial Illumina adaptor sequences and unique identifiers (barcodes) were attached by ligation to both the 5’ and 3’ ends of the PCR amplicons. A subsequent low-cycle PCR amplification step was used to complete the addition of full-length adaptor sequences. Paired-end sequencing (2 × 150 bp) of the resulting TruSeq-compatible paired-end Illumina libraries was performed on an Illumina MiSeq platform, using V2 chemistry.

### Targeting in HEK293T cells

HEK293T cells from one well of a 12-well plate were transfected with 0.4 μg AsCpf1 or SpCas9 plasmid, 0.8 μg targeting vector AAVS1-CAGGS-tdTomato using Fugene 6 transfection reagent (Promega). GFP and tdTomato double positive cells were sorted 3 days post transfection on a FACSAria (BD Biosciences). Sorted cells were plated at low density (2000 cells per 10 cm culture plate) and subjected to selection with 2 μg/ml puromycin until formation of colonies. Individual colonies were mechanically passaged to individual wells of 12-well plate for expansion.

For genotyping HEK293T clones with correct knockin, 0.5 μg of genomic DNA were used for PCR reaction with primer 495 and 489 (Supplemental Table 1) with Red taq (Sigma) with the following PCR condition: 95 **°**C 5 min, followed by 35 cycles of 95 **°**C 30 sec, 57 **°**C 30 sec, 72 **°**C 1min 40 sec, then 72 **°**C for 5 min. PCR products were resolved with 1% agarose gels. DNA from bands of expected size were extracted with gel purification kit (Omega Biotech), and subjected to Sanger sequencing with primer 495 (Supplemental Table 1).

### Targeting hPSCs

Undifferentiated human pluripotent stem cell line WIBR3 (Lengner et al., 2010) or iPS cells (Maetzel et al., 2014) was maintained in DMEM-F12 (Life Technologies cat# 11330-057) supplemented with 15% FBS, 5% KSR (Life Technologies cat# 10828-028), 4 ng/ml FGF (Life Technologies cat# PHG0261), 1X 2-Mercaptoethanol (Life Technologies cat #21985-023), 1X L-glutamine (Life Technologies cat# 25030-081), 1X MEM-NEAA (Life Technologies cat# 11140-050), 1X penicillin/streptomycin (Life Technologies cat# 15140-122). Cells were passaged with 1 mg/ml collagenase IV (StemCell Technologies cat# 07909) every 4-6 days on mitomycin C inactivated MEF feeders cultured in DMEM medium supplemented with 10% FBS, MEM-NEAA, 1X penicillin/streptomycin, and 1X L-glutamine. Undifferentiated human pluripotent stem cell line H1-OCT4-GFP (Zwaka and Thomson, 2003) was maintained in similar medium as above, except 20% KSR (Life Technologies cat# 10828-028) was used instead of 15% FBS plus 5% KSR.

Targeting of the pluripotent cell line was performed using previous methods (Ma et al., 2020). One day before electroporation, cells were cultured in the medium supplemented with Rho kinase (ROCK) inhibitor Y-27632 (Calbiochem). About 10 million hPSCs were dissociated into single cells with Accutase (Stemcell Technologies), filtered with a 40 μm cell strainer (Corning), washed with medium containing Y-27632, centrifuged, and resuspended in 800 µL PBS with 100 μg sg-*AAVS1* Cpf1 plasmid or 20 μg sg-AAVS1 SpCas9 plasmid, and 40 μg donor construct. The mixture was loaded to a 0.4-cm cuvette (BioRad), incubated on ice for 5 minutes before electroporation with a Gene Pulser Xcell System (Bio-Rad) with 1 pulse of 250 V, 500 μF. Then the cuvette was incubated on ice for 5 minutes before plating cells on 2 6-well plates of DR4 MEF (Tucker et al., 1997). Puromycin containing medium (0.5 μg/ml) was added 3-4 days after electroporation. Individual colonies were manually passaged onto MEF coated 12-well plate wells, expanded and tested with PCR and Southern blotting analysis.

### Southern blot

Southern blot was performed using *AAVS1* Southern blot probes as described previously(Ma et al., 2020). For *INS* targeting, internal probe was amplified with PCR with the following primers and firefly luciferase gene as template:

Forward: ATGGAAGACGCCAAAAACATAAAGAAAGGCCC

Reverse: CACGGCGATCTTTCCGCCCTT

External probe was amplified with PCR with the following primers and genomic DNA from H1 hPSCs:

Forward: GACTCCCCACTTCCTGCCCATCT

Reverse: TCTTCTCCCAGCCCCGTCCTCAC

### Immunofluorescence

Immunofluorescence analyses were performed according to previous methods (Li et al., 2014) with the following antibodies: mouse anti-OCT3/4, 1:100 (BD Transduction Laboratories, 611203); rabbit anti-NANOG, 1:100 (Thermo Fisher Scientific, PA1-097); mouse anti-TRA-1-60, 1: 100 (Life Technologies 411000); mouse anti-SSEA4 antibody, 1:100 (Thermo Fisher Scientific MA1-021, clone MC-813-70). Alexa 488 conjugated secondary antibodies (Life Technologies) were used at 1:400.

### Testing potential off-target effects with Sanger sequencing

PCR with primers highlighted in **Supplemental Table 1** were performed with 0.5 μg of genomic DNA from parental and targeted pluripotent stem cell clones. PCR products were resolved with 1% agarose gel followed by purification with gel purification kit (Omega Biotech), and Sanger sequencing with the sequencing primers listed in **Supplemental Table 1.**

### Differentiation of *INS* reporter hPSCs to β-like cells

The differentiation of β-like cells was based on a previous protocol (Ma et al., 2020).

### *In vitro* luciferase imaging

Undifferentiated H1-OCT4-GFP-INS:tdT-luciferase hPSC and the differentiated β-like cells were incubated with 15mg/mL D-luciferin for 1 min before imaging with an IVIS Spectrum system (PerkinElmer), with an exposure of 1 min and mid-bin setting. Imaging quantification were performed with Living Image (Version 4.5.4).

### Transplantation of hPSC-β-like cells, *in vivo* luciferase imaging, and human insulin measurements in mice blood

β-like cell clusters were transplanted using a 1 mL syringe with a 23-gauge butterfly needle (Terumo Corporation SV-23BLK) to the kidney capsules of 6-8 weeks old immunodeficient NOD.Cg-Prkdcscid Il2rgtm1Wjl/SzJ (NSG) mice (Jackson Laboratory). After 2-3 weeks, control and transplanted mice were intraperitoneally injected with D-luciferin (PerkinElmer 122799, 0.195 mg per gram weight), and imaged with an IVIS imager (PerkinElmer). Mid-bin setting of the imager and 5 min exposure were used. Living Image (Version 4.5.4) was used for imaging analyses. To measure human insulin levels in mouse blood, mouse blood samples were collected with heparin-coated blood collection tubes (Sarstedt Microvette; CB 300 LH). The collected blood samples were centrifuged at 3000 g for 3 min, and plasma samples were used for human insulin measurements with an Ultrasensitive Human Insulin ELISA kit (Mercodia) based on the manufacturer’s protocol. A linear model was used to generate standard curves.

### Teratoma assay

Human PSCs from two near confluent six-well plates were dissociated with collagenase, and resuspended in DMEM without serum. The cells were injected intramuscularly into NSG mice between 8-12 weeks of age using 1-mL syringes with 23-gauge needles. Formation of teratomas was monitored twice-weekly for about 2 months. Palpable tumors reaching between 7 and 10 mm in diameter, were dissected and fixed in buffer-neutralized formalin solution. Paraffin embedding and histological analysis (H&E staining) were performed with standard protocols. All animal experiments were performed in accordance with the protocols approved by the Animal Research Regulation Committee at the Whitehead Institute and guidelines from the Department of Comparative Medicine at Massachusetts Institute of Technology.

### Differentiation of neural precursors and neurons

Neuronal differentiation was performed based on published protocol (Muffat et al., 2016). Briefly, a total of 5 million dissociated WIBR3 or WIBR3-tdT cells were plated to one matrigel-coated 6-well plate well in NGD medium supplemented with 10 ng/ml FGF2, 1:500 insulin (Sigma), and 2.5 μM dorsomorphin for 14 days with daily medium change to keep medium from turning very yellow. Then NP cells were passaged, every 3 days, in NGD medium with 10 ng/ml FGF2, 1:500 insulin (Sigma). For neuronal differentiation, about 0.2 million NP cells were plated in each 12-well plate well in 1 ml of NGD medium. Cells were fixed after 4-6 weeks of differentiation with medium change every 2-3 days.

## ACKNOWLEDGMENTS

The authors wish to thank Feng Zhang’s laboratory for sharing pX458 and pY010 plasmid through Addgene, Huan Yang of Hongkui Deng’s laboratory for sharing *INS* targeting construct, Patrick Autissier, Eleanor Kincaid, and Hanna Aharonov of FACS core laboratory of Whitehead Institute for sorting cells, the Center for Computational and Integrative Biology (CCIB) at Massachusetts General Hospital for the use of the CCIB DNA Core Facility (Cambridge, MA), which provided NGS and data analysis service, members of the R.J. laboratory for advice and discussion. This study was supported by a generous gift from Liliana and Hillel Bachrach and in part by the NIH (RO1-CA084198) to R.J.

## Notes

### Competing Interest Statement

The authors have declared no competing interest.

