## Supplemental_figures for "Multiplex genome editing of human pluripotent stem cells using Cpf1"

**A**

```
CACCGTAATTTCTACTCTTGTAGATGTCTTCgaGAAGACtttttttt
|||||||||||||||||||||||||||||||||||||||||||||||
GTGGCATTAAAGATGAGAACATCTACAGAAGctCTTCTGaaaaaaaa
```

**B**

```
Oligo1 5’-GTAGATATCTGTCCCCTCCACCCCACAGT-3’
              ||||||||||||||||||||||||
Oligo2     3’-TATAGACAGGGGAGGTGGGGTGTCAAAAA-5’

            TTTN PAM          guide
Genomic  5’-TTTTATCTGTCCCCTCCACCCCACAGTGGGGCCACTAGGGACAG-3’
            |||||||||||||||||||||||||||||||||||||||||||
sequence 3’-AAAATAGACAGGGGAGGTGGGGTGTCACCCCGGTGATCCCTGTC-5’
```

**Supplemental Figure 1. Strategy to clone gene specific guide RNAs to the bicistronic AsCpf1 plasmids.**

(A) BbsI cloning sites of Cpf1 plasmids. Cyan denotes spacer sequence, gray denotes BbsI cognitive sites, and arrows denote cleavage sites upon BbsI digestion.

(B) An example of oligo design (top panel) for the targeted genomic sequence at the *AAVS1* locus (bottom panel). Cyan denotes the part of spacer sequence, yellow denotes gene specific sequence, red denotes ligation adaptor sequence, and magenta denotes PAM sequence in the genomic DNA.

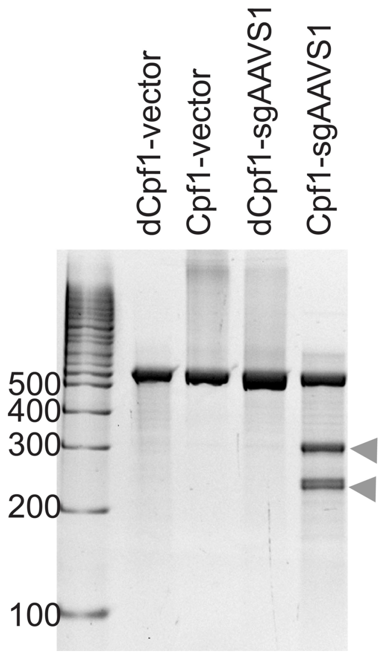


**Supplemental Figure 2. DNase activity is required for generating mutations by AsCpf1.**

T7EI assay with wild-type Cpf1, dead Cpf1 (dCpf1) that has a D908A mutation, or Cpf1, and dCpf1 with *AAVS1* guide.

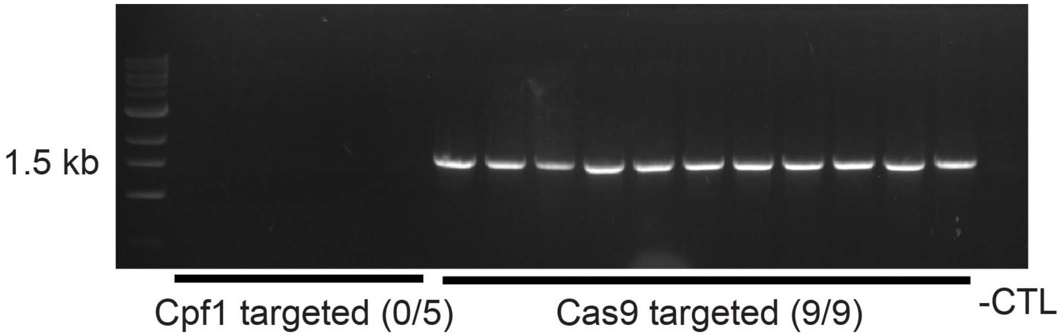


**Supplemental Figure 3. Comparison of CMV-SpCas9 and CMV-AsCpf1-mediated genome editing in hPSCs.**

Genomic DNA from H1 hPSCs electroporated with CMV-AsCpf1-*AAVS1*-sg plasmids (the first group of 5 samples) or with CMV-Cas9-*AAVS1*-sg plasmid (the second group of 9 samples) were subjected to PCR with primers spanning the homologues arm.

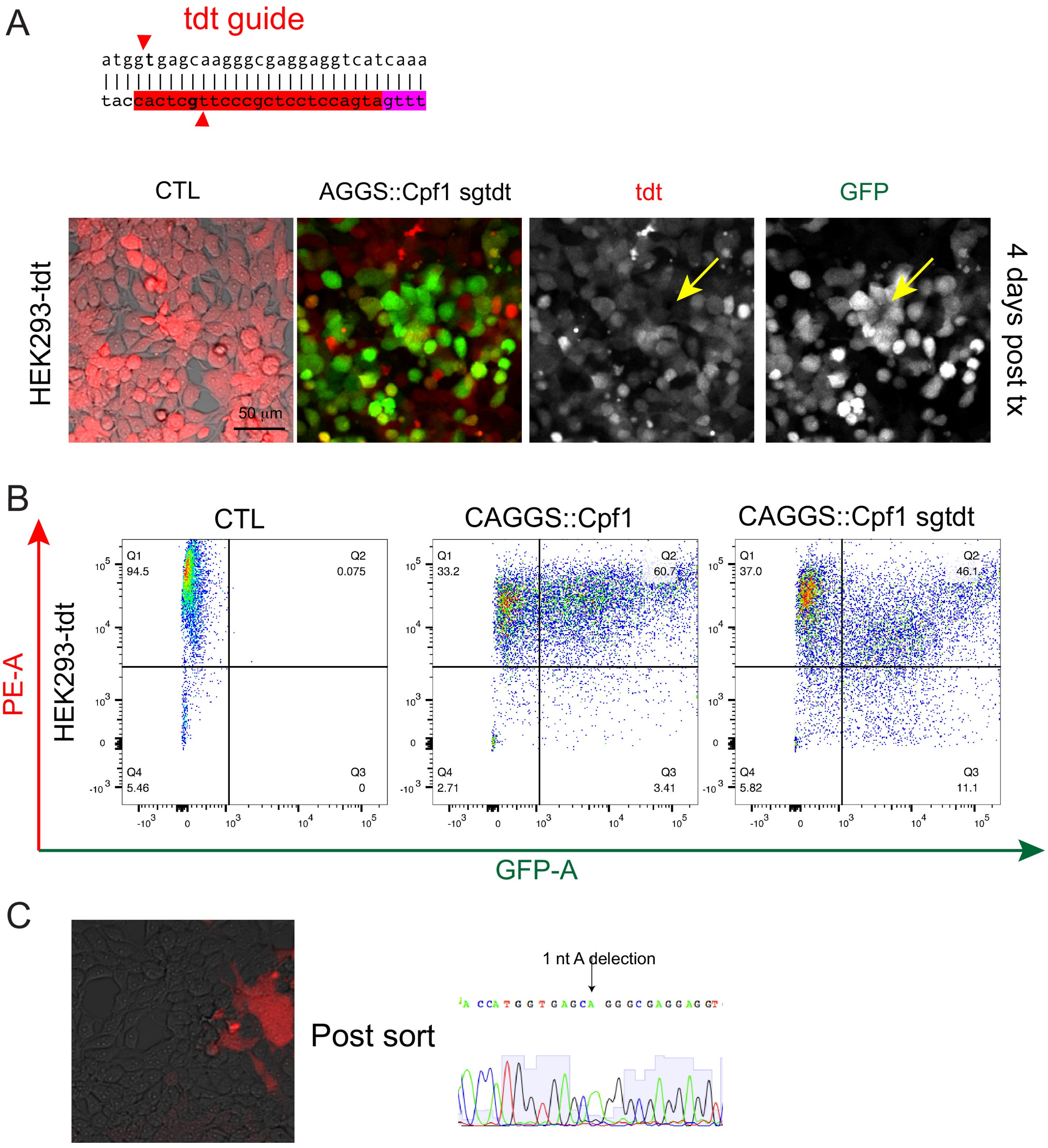


**Supplemental Figure 4. Disruption of tdTomato expression in HEK293T-tdT cells with CAGGS:Cpf1 construct that expresses a tdT guide RNA.**

(A) Top panel: a diagram showing the tdTomato guide. PAM sequence is highlighted in magenta, and guide sequences is in red. Expected double strand DNA breaks are indicated by the red arrowheads. Bottom panel, representative micrographs showing expressing of Cpf1 and the tdT guide RNA in HEK293T cells reduced tdTomato fluorescence signals.

(B) Flow cytometry analyses showing control HEK293T-tdT cells (left panel), HEK293T-tdT cells expressing AsCpf1 driven by CAGGS promoter (middle panel), and HEK293T-tdT cells expressing AsCpf1 and the tdT guide RNA.

(C) A representative micrograph showing cells plated after sorting for low expressing of tdTomato (left panel), and identification of 1 nucleotide deletion in the tdTomato coding sequences upon Cpf1-mediated gene editing.

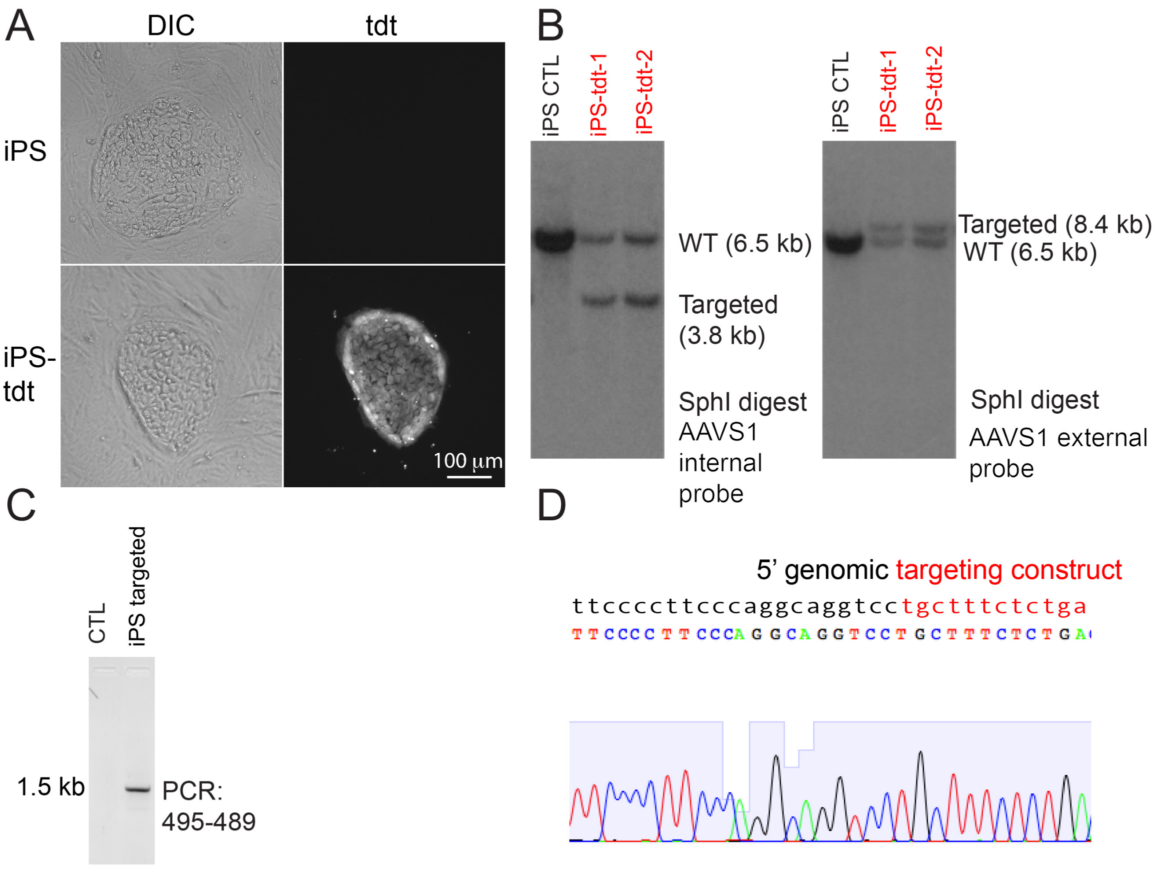


**Supplemental Figure 5. AsCpf1 mediated knockin in iPS cells.**

(A) Expression of tdTomato in targeted iPS cells.

(B) Southern blot confirming correct knockin of the *AAVS1* locus with internal probe (left panel) and external probe (right panel).

(C) PCR identification of knockin at the *AAVS1* locus in edited human iPS cells.

(D) Sequencing of PCR amplicon showing correct knockin.

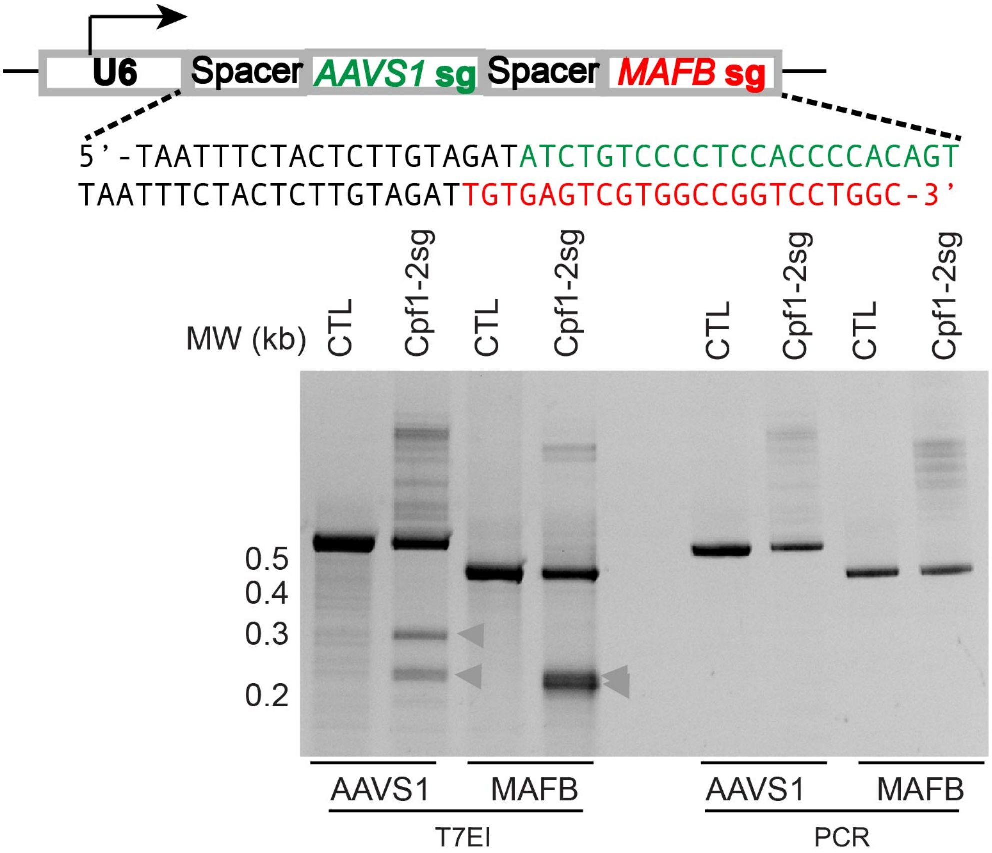


**Supplemental Figure 6. Simultaneous genome editing of *AAVS1* and *MAFB* loci in HEK293T cells with a guide RNA array.**

Top panel: design of an *AAVS1* and *MAFB* targeting spacer guide RNA array driven by a U6 promoter. Bottom panel: T7EI assay showing mutation formation at both targeted loci in HEK293T cells expressing AsCpf1 and the spacer guide RNA array.

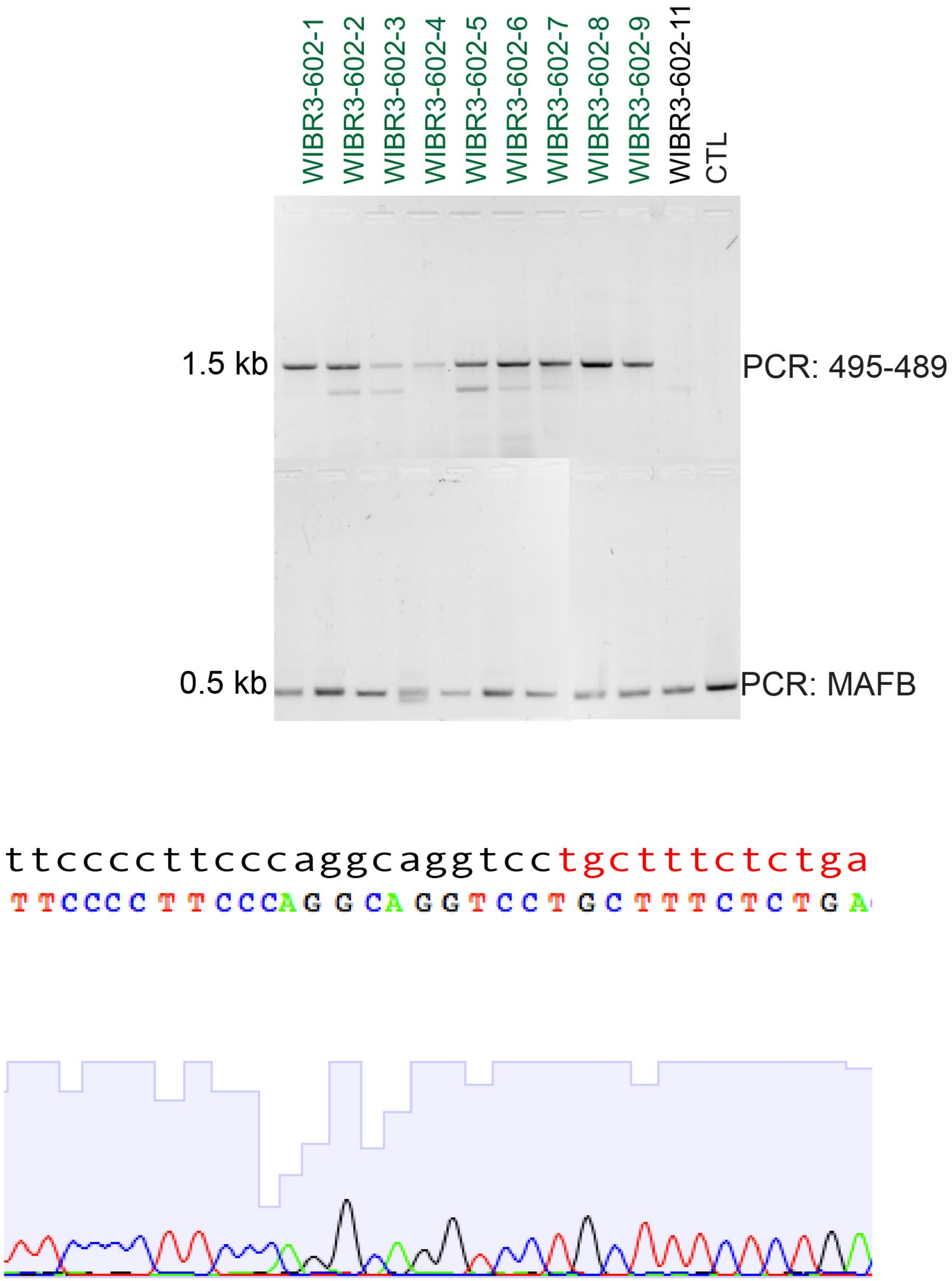


**Supplemental Figure 7. PCR-based characterization of WIBR3 cells targeted with the CAGGS-Cpf1-2A-GFP-AAVS1-MAFB construct.**

PCR analyses of knockin at the *AAVS1* locus (top panel) and the *MAFB* locus (middle panel). Clones showing correct knockin are labeled in green. Representative sequencing results of PCR amplicons to verify correct knockin at *AAVS1* locus (bottom panel).

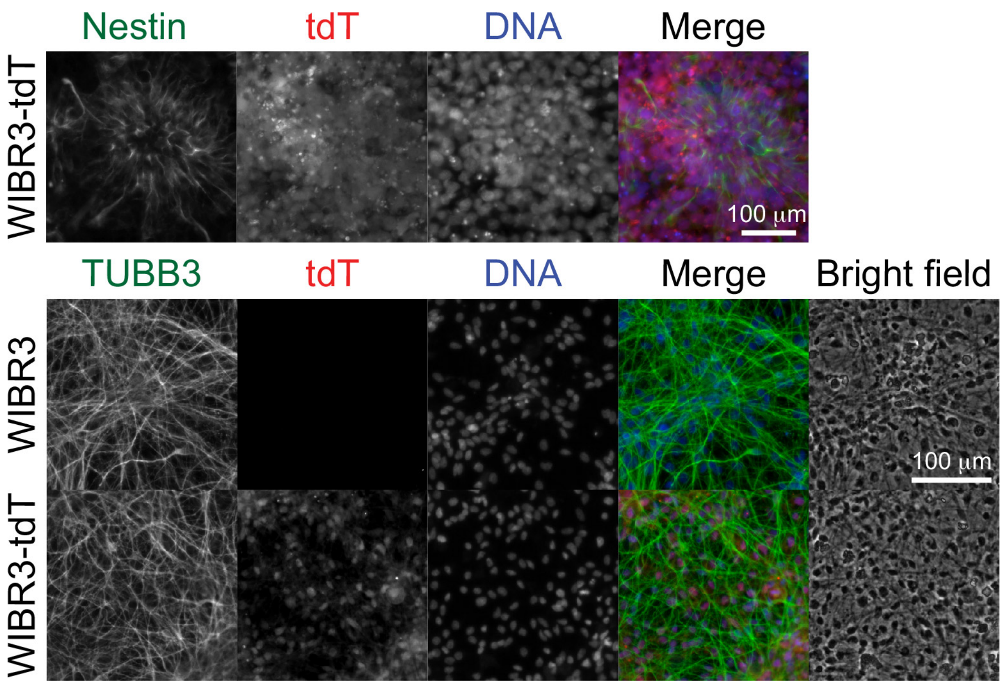


**Supplemental Figure 8**. **Differentiation of Cpf1 edited WIBR3-tdT cells to neural precursor cells and neurons.**

Top panel: WIBR3-tdT cells were differentiated to neural precursor (NP) cells for 14 days. Then the NP cells were passaged, fixed and stained with anti-Nestin antibody (1:100, SC-23927). The formation of tdT-expressing (red) neural rosette structures (green) was shown.

Bottom panel: The WIBR3-tdT differentiated NP cells were differentiated toward neurons with growth factor withdrawal for four weeks. Neurite structures were shown by anti-TUBB3 (1:1000) staining.

**Supplemental Table 1. List of primers**

| Primers | Sequence | Note |
|---|---|---|
| 439-1 | GTAGATATCTGTCCCCTCCACCCCACAGT | Primers for p-74 (p-63-AAVS1 23 nt) |
| 440-1 | AAAAACTGTGGGGTGGAGGGGACAGATAT | |
| 445 | GTAGATTGTGAGTCGTGGCCGGTCCTGGC | Primers for p-75 (p-63-*MAFB* 23 nt) |
| 446 | AAAAGCCAGGACCGGCCACGACTCACAAT | |
| 481 | GTAGATCGTCGCCGTCCAGCTCGACCAGG | Primers for p-77 (p-63 *GFP* 23 nt) |
| 482 | AAAACCTGGTCGAGCTGGACGGCGACGAT | |
| 14 | CACCGGAAAGAACTCGGGAGAGGAG | Primers for p-6 (458-*MAFB*) |
| 15 | AAACCTCCTCTCCCGAGTTCTTTCC | |
| 8 | CACCGGCTGTCCTGAAGTGGACATA | Primers for p-8 (458-*AAVS1*) |
| 9 | AAACTATGTCCACTTCAGGACAGCC | |
| 483 | TTTGCTGCCTCCAGGGATCCTGTGTCC | NGS primer for *AAVS1* locus |
| 484 | GACCCAATATCAGGAGACTAGGAAGGAGG | |
| 425 | GGCAAAGAATTCCGCCACCATG | Surveyor assay primers for GFP |
| 92 | CTTGTACAGCTCGTCCATGCC | |
| 630 | GCAACGTGCTGGTTATTGTGCTG | Testing mutation in H1-tdt cells, amplicon 158 bp |
| 631 | TCGCCCTCGCCCTCGATCTCGAA | |
| 10 | CCCCTTACCTCTCTAGTCTGTGC | Surveyor assay primers for *AAVS1* |
| 11 | CTCAGGTTCTGGGAGAGGGTAG | |

| | | |
|---|---|---|
| 18 | GGGGCTACGCCCAGTCTTGCAGGTATAAACG | Surveyor assay primers for *MAFB* |
| 19 | TCTCTCTCTCCGGCTCTGCTCGAGTCTAGGAGG | |
| 417-1 | TGGGGCAGGTGGAGCTGGGCGG | Surveyor assay primers for *INS* |
| 418-1 | ACCACCCCTGGCCCCTCAGAGACC | |
| 495 | TCTCTCTCCTGAGTCCGGACC | For detection of knockin of puro resistance gene at the *AAVS1* locus |
| 489 | ACTGAGCTCTCAGGCACCGGGCTTGCGG | |
| aavs1 off 1 F | CTGCTGAACACTCAGCATCTGCC | Sequencing primer |
| aavs1 off 1 R | CAGAGGAGCGAGTGGAGCAGACAG | |
| aavs1 off 2 F | ATTGCAATATCCTCCTATTAGCC | Sequencing primer |
| aavs1 off 2 R | CACTAGAGTCACCCTATGGCTCCC | |
| aavs1 off 3 F | CATGGGACAAGTTGATAGCTAAG | Sequencing primer |
| aavs1 off 3 R | GAAGTTACTCTGAAACGTATAGCAC | |
| aavs1 off 4 F | GCTGCTCCTGGATTTAGCAAAC | Sequencing primer |
| aavs1 off 4 R | GCCCAGCCCAACACTTTTGGTC | |
| aavs1 off 5 F | GTGTGCTAGTATCATTGCAAAAG | Sequencing primer |
| aavs1 off 5 R | CACTATTGCGTTCTCTCATTTCTC | |
| MAFB off 1 F | TCTGAGGTCCGTCTCACACACTG | |
| MAFB off 1 R | CACTGTTCAAAGAGTTTGAACATTCC | Sequencing primer |
| MAFB off 1 F | CTCCTGACTATTGCAGTTGCTGGTCACC | Sequencing primer |
| MAFB off 1 R | AACAGAGGAGCGAGTGGAGC | |
| INS off 1 F | CGGAGTCTCACTTTGTTGCCATG | Sequencing primer |
| INS off 1 R | TGGTTAGAACTTCCTGCCCACAG | |

| | | |
|---|---|---|
| Luciferase probe forward | ATGGAAGACGCCAAAAACATAAAGAAAGGCCC | *INS* targeting internal probe |
| Luciferase probe reverse | CACGGCGATCTTTCCGCCCTT | |
| 665 | GACTCCCCACTTCCTGCCCATCT | *INS* external probe forward |
| 666 | TCTTCTCCCAGCCCCGTCCTCAC | *INS* external probe reverse |
| 662 | GGCCTTTGGTGCAGTGACCAGAGTGTCAGG | *INS* knockin genotyping primers |
| 663 | GATTCTCCTCGACGTCACCGCATGTTAGC | |

**Supplemental Table 2 List of constructs**

| Construct | Note |
|---|---|
| p-63 | CMV:AsCpf1-2A-GFP, guide RNA can be cloned into BbsI sites |
| p-74 | p-63-AAVS1 guide |
| p-77 | p-63-GFP guide |
| p-75 | p-63-MAFB guide |
| p-76 | p-63-INS guide |
| p-6 | pX458-MAFB guide |
| p-8 | pX458-AAVS1 guide |
| p-93 | p-63-tdTomato guide |
| Addgene 159281 | CAGGS:AsCpf1-2A-GFP, guide RNA can be cloned into BbsI sites |
| p-70-AAVS1 | CAGGS:AsCpf1-2A-GFP-AAVS1 guide |
| p-70-tdT | CAGGS:AsCpf1-2A-GFP-tdTomato guide |
| Addgene 159283 | CAGGS:AsCpf1-2A-GFP-INS guide |
| p-70-602 | CAGGS:AsCpf1-2A-GFP with a guide RNA array targeting both AAVS1 and MAFB |
| p-69 | CAGGS:dAsCpf1(D908A)-2A-GFP, guide RNA can be cloned into BbsI sites |
| p-69-AAVS1 | pHM-wi-69-AAVS1 guide |
| p-61 | AAVS1-tdTomato targeting vector |
| Addgene 159348 | INS luciferase-tdTomato targeting |
